# PEAS: Detection of Clustered Differences in Genomic Data

**DOI:** 10.1101/2025.03.31.646436

**Authors:** Dylan Derik Skola, Christopher Glass

## Abstract

**Motivation:** Studies of genetically-distinct individuals have shown that differences in marks of transcriptional regulation such as chromatin accesibility, transcription factor binding and histone modifications are often proximally clustered along the genome. These proximal clusters, which have been labeled as *cis*-regulatory domains (CRDs), are thought to reflect topological features of the genome and may demarcate functional units linking genetic variation to transcriptional regulation. The problem of distinguishing CRDs from background variation is computationally difficult and current methods rely on greedy approaches with ad-hoc parameters and do not provide an assessment of statistical significance, an important consideration for investigating CRDs in small sample cohorts.

**Results:** We developed a software package, *PEAS* (Proximal Enrichment by Approximated Sampling), to identify CRDs from a small number of samples (as few as two distinct genetic backgrounds) using a robust statistical approach. *PEAS* uses methods for efficient and accurate estimation of empirical distributions to quantify the significance of enriched regions, followed by a dynamic programming algorithm to identify the minimum likelihood set of non-overlapping enriched regions. We used it to identify clusters of proximally-enriched differences in the histone mark H3K27ac between two mouse strains as well as proximally-enriched regions of correlation in this mark across five mouse strains. We find that differences in histone acetylation between two mouse strains form signficant clusters that overlap closely with differences in the first principal component of their Hi-C correlation matrices.

**Availability:** *PEAS* is written in Python and is available at https://pypi.org/project/PEAS/. Methods for approximating empirical distributions are implemented in C and Python and are available at https://pypi.org/project/empdist/.

## Introduction

Studies of chromatin accessibility using assay for transposase-accessible chromatin using sequencing (ATAC-seq) (2) as well as assays of transcription factor binding and histone and histone tail modifications using chromatin immunoprecipitation followed by sequencing (ChIP-seq) (9) in the presence of genomic variation have shown that inter-individual differences in the magnitude of these marks often form proximal clusters along the genome. Correlating ATAC-seq profiles of activated T-cells from 105 human individuals demonstrated patterns of coordinated variation at multiple scales that closely matched the correlation pattern of Hi-C interactions for those regions (7). Profiling of lymphoblastoid cell lines (LCLs) from hundreds of human individuals revealed modules of coordinated variation that preferentially clustered within chromatin contact domains and that were associated with sequence variation in TF-bound regions of *cis*-regulatory elements (18; 4). A smaller-scale approach using bone-marrow-derived macrophages from five strains of laboratory mice revealed modules of coordinated regulation across multiple data types (11). Several of these studies have sought to computationally determine the boundaries of these modules of genetic variation, known variously as variable chromatin modules (VCMs) (18), cis-coaccessibility networks (CCANs) (16), or cis-regulatory domains (CRDs) (4; 11).

However, current methods used to identify CRDs suffer from a lack of robustness and statistical rigor, which limits their interpretability and applicability. The general approach is to first compute, in a pairwise fashion, some measure of correlation of the activation (as measured by normalized ATAC-seq or ChIP-seq tag counts over multiple samples) between genomic loci. Given these correlations between individual loci, the computational challenge is to define sets of loci that exhibit mutually correlated behavior while excluding loci that vary in an uncoordinated fashion. To date this has been done by applying a threshold on the mean correlation of loci enclosed by the CRD. For example, after computing the pairwise correlation of peaks with their neighbors in a 250-peak window, Delaneau and colleagues perform hierarchical clustering on the rows of the correlation matrix and cut the dendrogram into CRDs using the following heuristics:

1. The mean pairwise (absolute) correlation of peaks in the CRD must be more than twice the mean correlation of all peaks in the chromosome
2. The mean correlation of the peaks at the boundaries of the CRDs with the internal CRD peaks must be greater than twice the mean correlation of all peaks with the the first and last peaks on the chromosome.
3. The CRD must enclose at least two non-overlapping merged chromatin features

**Figure 3.1:**
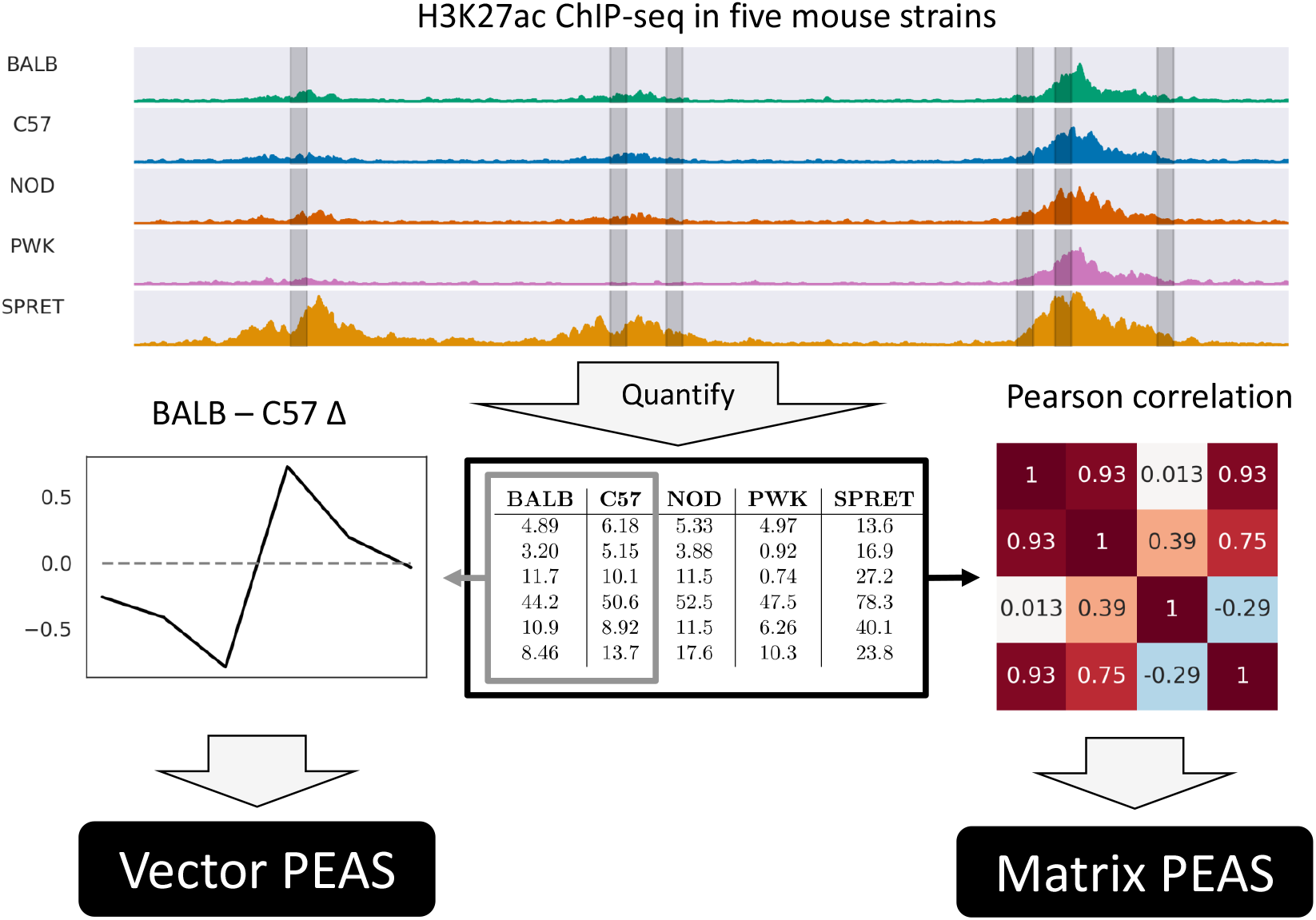
Workflow for identifying CRDs with PEAS. First, ATAC-seq or ChIP-seq tags are quantified at non-overlapping loci across multiple samples, which could represent different genetic background, stimulation conditions, tissue types, developmental trajectory, etc. After normalization, two samples can be compared to produce a vector of differences which is analyzed using the Vector PEAS approach to find statistically-significant regions of coordinated difference. Alternatively, the pairwise correlations of the per-sample magnitudes can be computed and the resulting matrix analyzed using the Matrix PEAS approach to find enriched or depleted regions of correlation.

Such a method is obviously sensitive to the particular distribution of correlation values across the chromosome. Statistically, the first rule is equivalent to setting a *p*-value threshold of 1 − *F* (2*µ*). However, the value of *F* (2*µ*) and therefore the equivalent *p*-value will vary greatly depending on the underlying distribution. If all the peaks across that chromosome exhibit inflated correlation as a result of biological effects or technical artifacts then the proportion of regions exceeding 2*µ* will be quite different than if correlations are depressed because of technical noise. Another obvious consequences is that, if the mean correlation across all peaks is greater than 0.5 then no regions will be defined as CRDs. Likewise the arbitrary use of the first and last peak on the chromosome to define the null expectation of edge correlation is sensitive to the particular sequence of peaks on that chromosome.

In Pliner et al. (16), the authors apply a similar approach to single-cell ATAC-seq data that varies along a developmental trajectory, first smoothing individual cells into related clusters, then using graphical LASSO to compute a regularized correlation of the accessibility of sites within 500 kb. After hierarchical clustering, CCANs are identified as clusters whose mean correlation exceeds a user-defined threshold. Although this avoids the pitfalls of using 2*µ*, as above, it introduces an additional free parameter that is sensitive to data quality (e.g. random technical noise will reduce observed correlations and systematic technical error will increase it).

While the use of hierarchical clustering in previous approaches allows for the identification of non-contiguous CRDs, the data explored with these methods have consisted of relatively large sample cohorts derived from dozens or hundreds of genetically distinct individuals. In order to power investigations of this phenomena in difficult-to-obtain human primary cells, methods are required that can reliably and robustly identify CRDs in smaller sample sets.

### Approach

In this article we present a method, proximal enrichment by approximated sampling (PEAS), that allows researchers to quantify the significance of CRDs under a simple null model that requires no assumptions about the distribution of individual relationships (and therefore places less stringent demands on data normalization). Applying a *p*-value cutoff is a familiar process and the meaning of a given threshold is data-independent and easily comprehended. Accurate computations of *p*-values also allow the use of methods for correcting for multiple hypothesis testing - an important consideration in genome-wide searches. By including the constraint that CRDs must comprise linearly-contiguous regions we reduce the potential for false positives which enables discovery with small sample sizes.

The method can be applied to the results of an assay such as ATAC-seq or ChIP-seq that generates quantitative data at various locations across the genome. After performing the same assay in different contexts (for example, on cells from genetically-distinct individuals or in the presence and absence of some stimulus) and quantifying the results at the same loci, the resulting data table can be processed with PEAS in two ways (figure 3.1): first, to identify CRDs using differences between a single pair of samples in an approach we refer to as Vector PEAS (figure 3.3), or using correlations between loci across three or more samples in an approach refered to as Matrix PEAS (figure 3.4).

### Data

We downloaded two replicates each of ATAC-seq and ChIP-seq for acetylation on histone 3 lysine 27 (H3K27ac) sequencing reads for unstimulated bone-marrow-derived macrophages (BMDMs) from five strains of mice (C57, BALB, NOD, PWK, and SPRET) that had been aligned and lifted over to mm10 coordinates (11). We generated tag directories using HOMER with the parameter -tbp 1 (to eliminate PCR duplicates), pooled the replicates and called 200 bp peaks on the pooled ATAC-seq data using HOMER (8). We used the merge function of bedtools (17) to merge the coordinates of overlapping peaks across the strains. Finally, we used the annotatePeaks command of HOMER to count the H3K27ac ChIP-seq reads in a 400 bp window around the center of each merged ATAC-seq peak. The read counts were normalized to give each strain a total of 10 million reads in peaks.

To compute the SPRET-C57 Δ vector we added a pseudocount of 5 to the normalized tag counts, took the base-2 logarithm and subtracted the logarithm of the pseudocount (to ensure that tag counts of zero had a value of zero after normalization. We then subtracted the C57 values from the SPRET values to produce the Δ vector. To compute the five-strain correlation matrix we computed the pairwise Pearson correlation of the normalized (but not log-transformed) tag count vectors at each locus.

From the same study we also obtained the Hi-C data generated in SPRET and C57 BMDMs (11) and used HOMER’s AnalyzeHiC command with a bin-size of 20 kbp and spacing of 10 kbp (“super-resolution mode”) to generate normalized count matrices for each chromosome in each strain. We then computed the Pearson correlation of the columns of this matrix to generate a Hi-C correlation matrix for each chromosome in each strain. The first principle component of each correlation matrix was computed and oriented so that positive values correspond to ‘A’ compartments (5).

**Figure 3.2:**
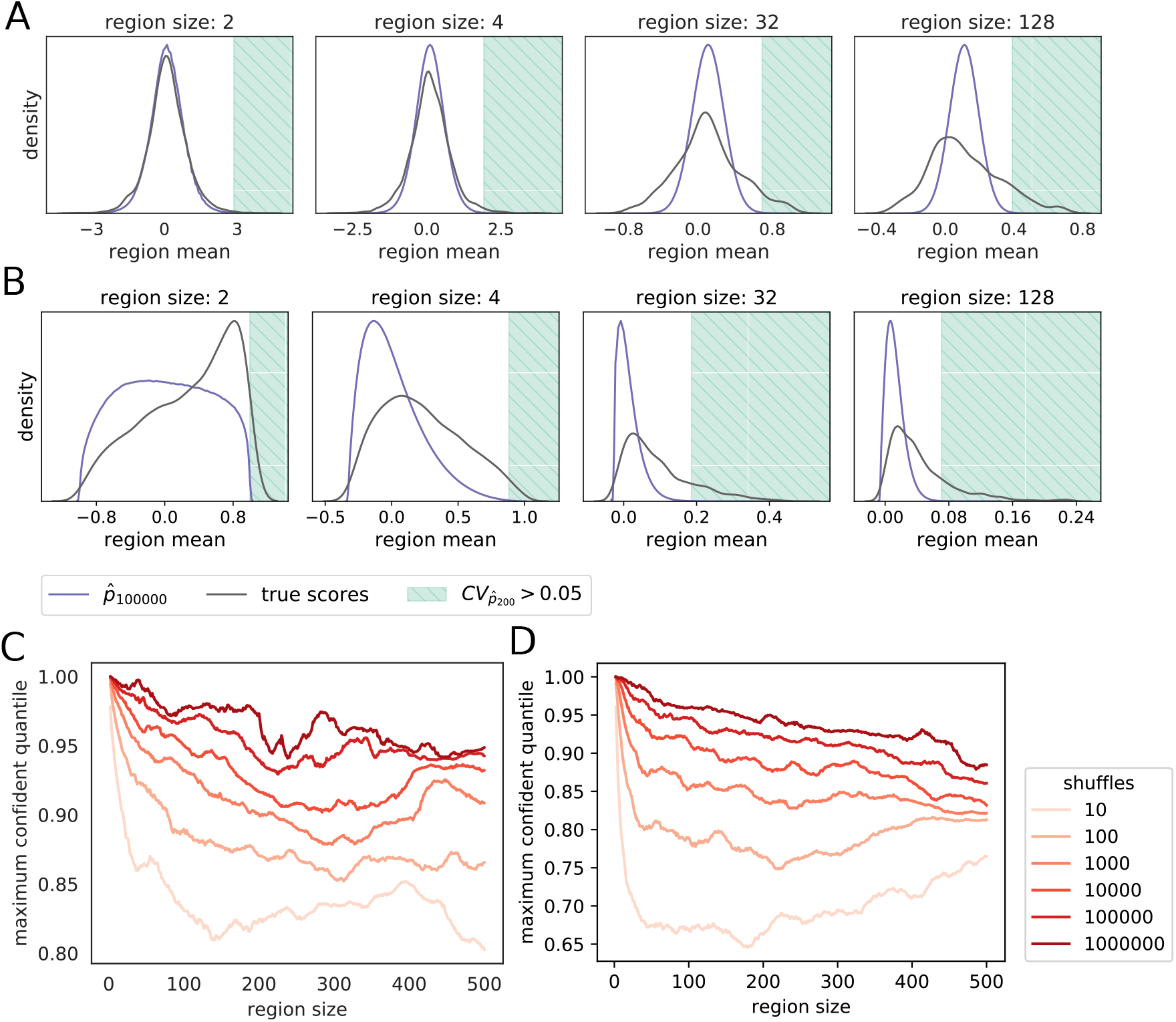
(A) Distributions of means of the chromosome 19 H3K27ac ChIP-seq SPRET-C57 Δ vector for regions of the indicated size along chromosome 19 (“true scores”), together with the ground truth null distributions. Cross-hatched area indicates region of right tail where *p*-values estimated using the 1,000 permutation will have unacceptable variance. (B) The same distributions computed for all means of pairwise Pearson correlations of chromosome 19 inter-strain H3K27ac ChIP-seq tag counts. (C,D) The maximum quantile of the observed region scores whose significance can be computed using null distributions derived from the indicated number of shuffles of chromosome 19 for the H3K27ac ChIP-seq SPRET-C57 Δ vector (C) and pairwise Pearson correlations of inter-strain H3K27ac ChIP-seq tag counts (D).

## Methods

### Vector PEAS finds significant regions of extreme values in a sequence of observations

Consider an ordered sequence of *n* observations 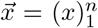, in this example: differences in normalized H3K27ac ChIP-seq tag counts between SPRET and C57 strains of mice along chromosome 19. We want to identify subsequences 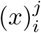 of this sequence that consist of adjacent genomic loci that exhibit scores that are unlikely under the null hypothesis that the order of the observations is not important.

The first step is to compute an upper-triangular matrix *S* containing the scores for all possible subsequences of 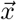.

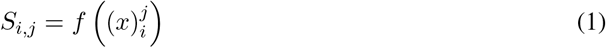

where *f* is a function that generates a scalar summary statistic for a given subsequence.

Possible scoring functions include the mean, minimum and maximum. These can be computed using dynamic programming in *𝒪* (*n*^2^) time. For the max function (the min is analogous) the following recurrence relation is used:

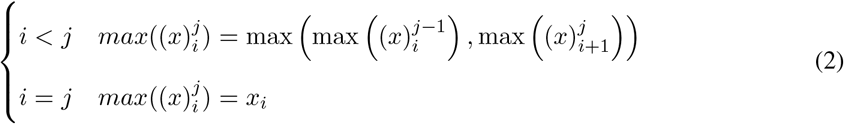

To compute the means of all possible subsequences we first compute their sums using the recurrence relation:

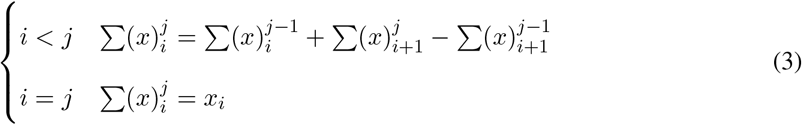

then:

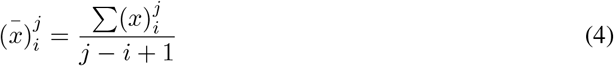

**Figure 3.3:**
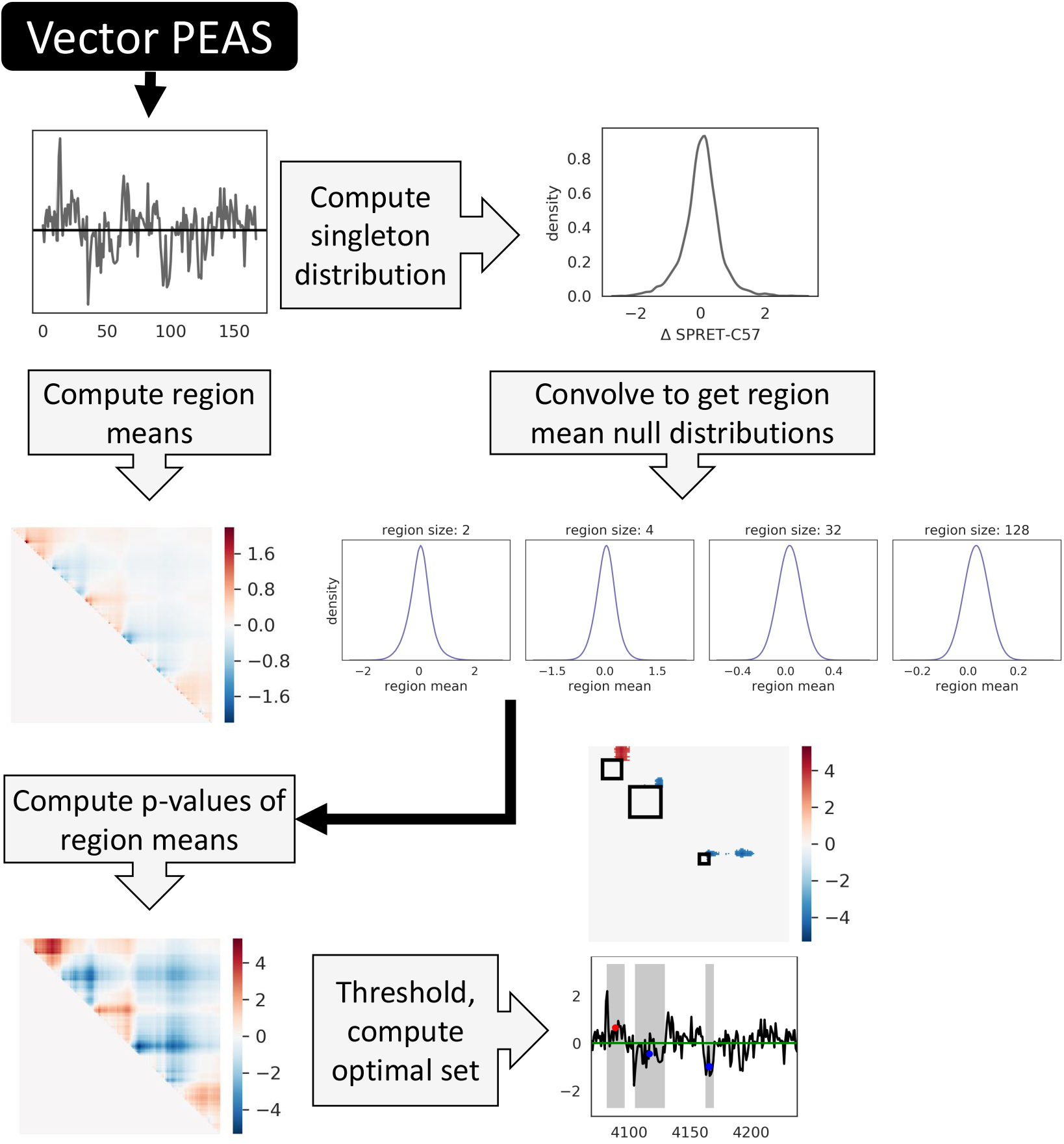
Workflow for vector PEAS. The means of all contiguous regions of the input Δ vector are calculated to obtain a score matrix. A histogram approximation of the distribution of vector elements is repeatedly convolved with itself to generate null distributions that are then used to compute the p-value of each region mean. Finally, a dynamic programming algorithm is used to find the minimum likelihood set of non-overlapping regions that meet the threshold criteria.

Although multiple scoring functions are implemented in `empdist`, for simplicity we will consider only the mean scoring function for the remainder of this manuscript.

Once we have computed the score matrix *S*, the next step is to compute the significance of the score of each region under the null hypothesis. Rather than make assumptions regarding the distribution of differences between sequence elements, we test only the significance of their local enrichment by using the scores of shuffled sequences as a null distribution. Since more extreme scores will occur more frequently by chance in smaller subsequences, a separate null distribution must be computed for each subsequence length.

To do so, we adopt an empirical approach to computing the nulls that relies on the fact that the distribution of the sum of two independent random variables is the convolution of the distributions of the original variables. In the case of empirical distributions of continuous random variables, we can approximate their probability distribution by generating frequency histograms of the data. For a histogram approximation with *b* bins, the cumulative frequency at each bin can be pre-computed, allowing accurate estimates of empirical *p*-values to be returned in the 𝒪(log *b*) time required to find the appropriate bin rather than directly from the data in 𝒪(log *n*) time.

### Expectation of arithmetic operations on independent random variables

We can compute the distribution of the sum *Z* of two random variables *X* and *Y* by convolution of their histogram approximations:

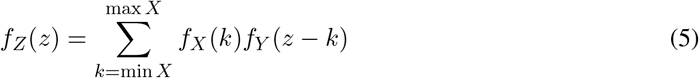

**Figure 3.4:**
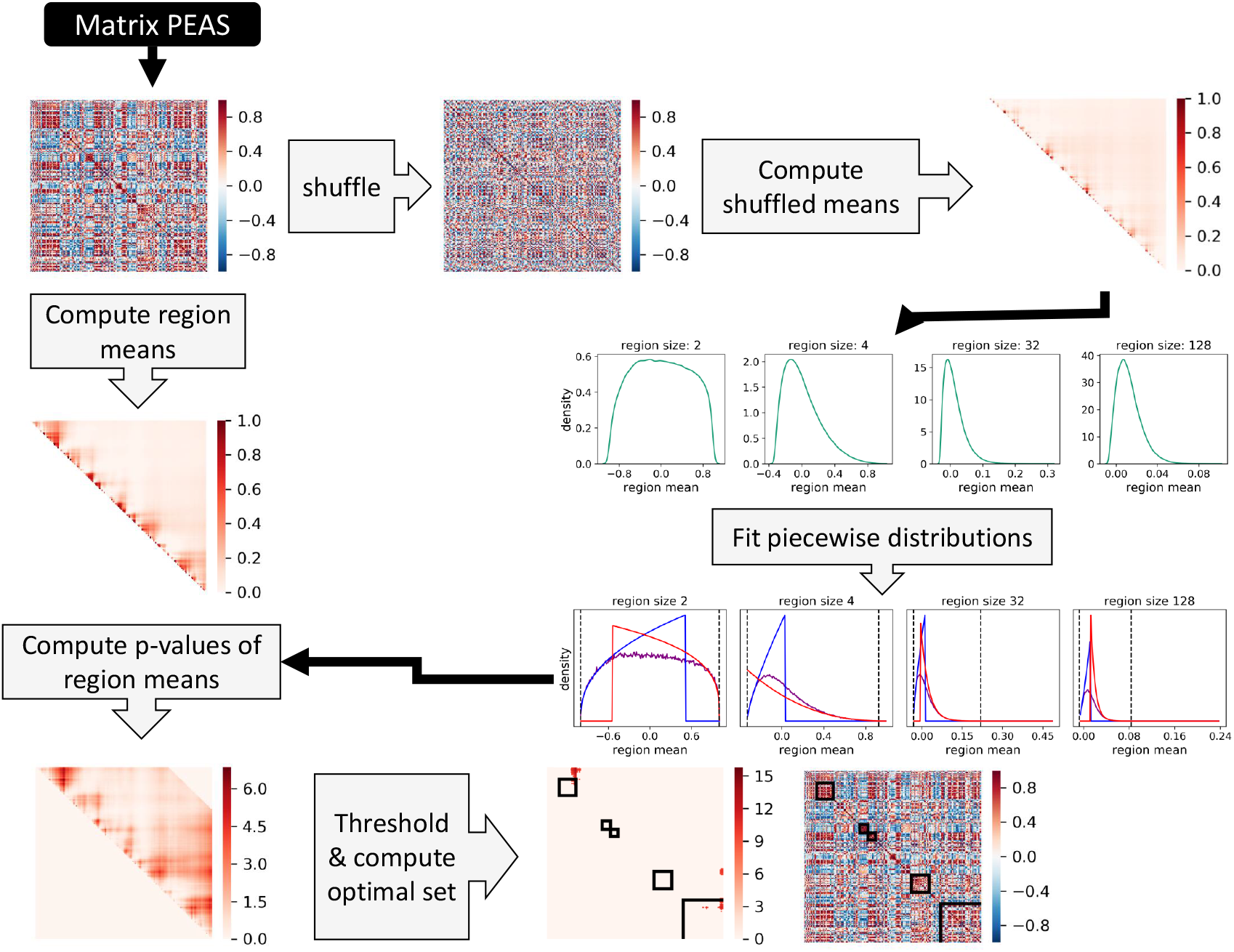
Workflow for matrix PEAS. The means of all contiguous regions are calculated to obtain a score matrix. The correlation matrix is repeatedly shuffled and the resulting region means produce an empirical null distribution for each region size. A piecewise null distribution consisting of a histogram center and generalized Pareto tails is fit to each empirical null distribution and used to compute the p-value of each region mean in the unshuffled data. Finally, a dynamic programming algorithm is used to find the minimum likelihood set of non-overlapping regions that meet the threshold criteria.

The support of the resulting histogram then becomes

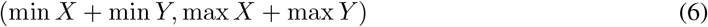

Simple manipulations of the support of each distribution allow implementation of the negation, min and max operators.

Let 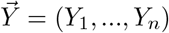 represent a vector of random variables whose *i*th component represents the sum of *i* elements drawn randomly without replacement from *x. Y*_1_ is then the empirical distribution of *x* and by generating a histogram approximation to that distribution, the distribution of *Y*_*i*_ can be predicted by convolution of *f* (*Y*_*i*−1_) with *f* (*Y*_1_). The distribution of the mean of a random sample of size *r* is then 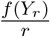.

Although binning introduces some quantization error, this method retains accuracy even after hundreds of repeated convolutions, in part because the number of bins is allowed to grow throughout the convolution, and the increasing resolution mitigates the propagated error. If the number of bins becomes computationally intractable, the histograms can be resampled to a lower resolution at the expense of some accuracy.

### Piecewise hybrid approximation of empirical distributions (Matrix PEAS)

We next consider the related problem of finding contiguous regions of observations whose elements are enriched or depleted in some pairwise relationship relative to expectation under a shuffled null distribution. In the case of CRDs this relationship is the Pearson correlation between sample-wise data vectors across genomic loci but the problem and solution are general for any matrix of pairwise relationships among ordered observations.

Let *X* be a matrix of observations such that *X*_*i*,*j*_ represents the relationship between ordered loci *i* and *j*. As we do for the vector method, we first compute a score matrix *S* efficiently using modified versions of the recurrence relations given in equations 2 and 3 that incorporate an additional term for the airwise relations. In order to account for situations where adjacent loci may have artificially-inflated correlations (for example in some types of Hi-C correlation) we allow the user to specify the minimum diagonal *k* (equivalent to a region size of *k* + 1) to be included in the region scores. So, for example the sum matrix is computed with the recurrence relation:

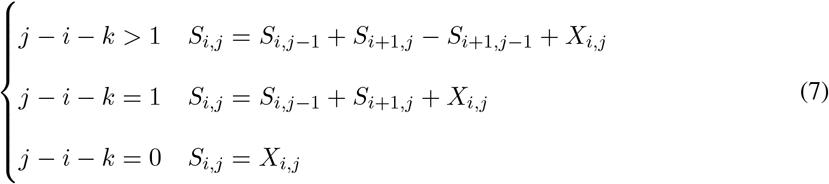

**Figure 3.5:**
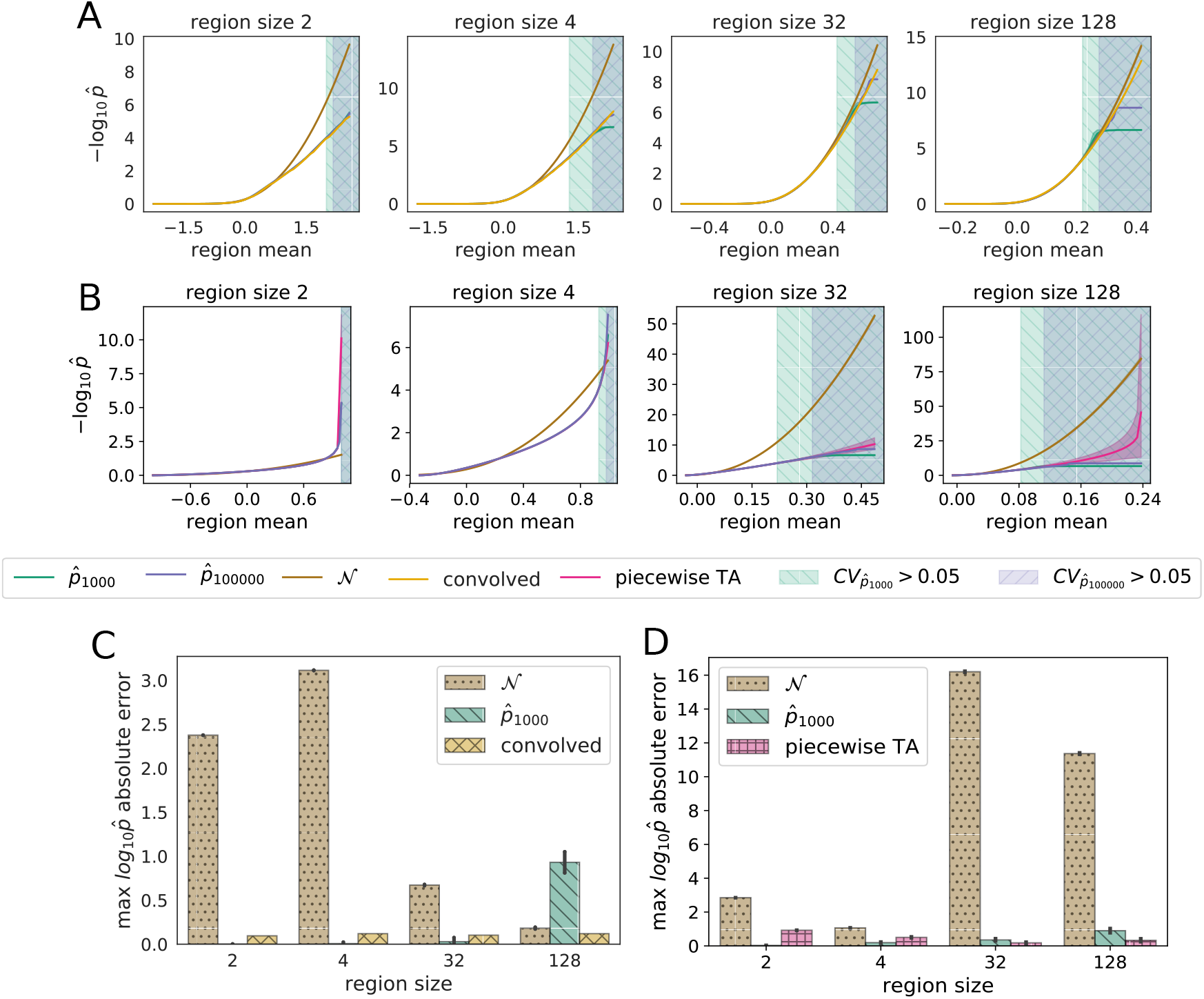
(A,B) Negative log_10_ 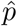 values as a function of region score for various size regions of the chromosome 19 H3K27ac ChIP-seq SPRET-C57 Δ vector (A) and pairwise Pearson correlations of chromosome 19 inter-strain H3K27ac ChIP-seq tag counts (B). Estimations are computed using 32 independently-generated 1,000 shuffle empirical distributions 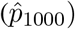, a theoretical normal distribution fit to the 1,000 shuffle data, a 100,000 shuffle empirical distribution 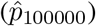 as well as the two methods presented in this article: estimation using convolution of histograms (convolved, (A) only) and using a piecewise tail approximation (piecewise TA, (B) only). Regions of the score distribution that cannot be confidently predicted at *CV*_*p*_ *<* 0.05 using the 100,000 shuffle data are shaded in blue and those that cannot be confidently predicted at *CV*_*p*_ *<* 0.05 using the 1,000 shuffle data are shaded in green. Estimations deriving from the 32 x 1,000 shuffle data are plotted as the range between the first and third quartiles. (C,D) Maximum deviation of estimated log 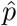 values using the indicated method from those estimated using the 100,000 shuffle data set for the chromosome 19 H3K27ac ChIP-seq SPRET-C57 Δ vector (C) and pairwise Pearson correlations of chromosome 19 inter-strain H3K27ac ChIP-seq tag counts (D).

Since most pairwise metrics of interest will be at least partially transitive (*e*.*g*. if region *a* is correlated with region *b* and *b* is correlated with region *c* then region *a* is more likely to be correlated with *c*), the assumption of independence that enabled the histogram convolution approach to the one-dimensional problem no longer holds. A common approach to such problems is to generate empirical null distributions by permutation testing, where a randomized null model is run repeatedly to generate data to which the observed value is compared to estimate the likelihood of the observation under the null hypothesis.

The minimum *p*-value that can be computed by empirical estimation with *n* samples is 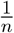. However, the variance in the estimates of *p*-values at the limit may be unacceptably high. Since an empirical 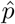 estimates *p* by the the proportion of samples more extreme than the queried data point, we can use the standard error of the proportion to compute the expected standard deviation in the estimates of a given value of *p*:

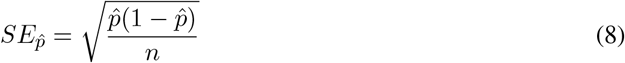

If we set an acceptable threshold for the coefficient of variation of 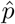 as 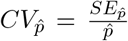 then the minimum number of samples needed to achieve an expected 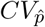 is:

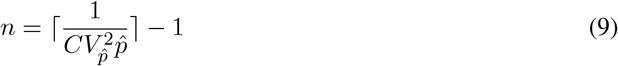

Similarly, we can compute the minimum *p*-value that can be estimated at a given 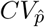 threshold:

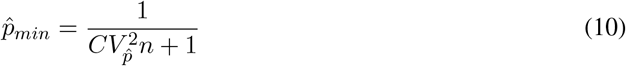

We can obtain *n* unique null samples either by sampling configurations without replacement or, for *d* samples with replacement the expected number of unique samples is:

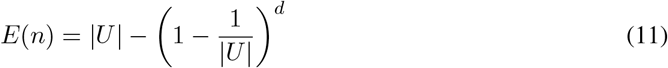

where *U* is the universe of possible samples. For the case of a contiguous region of size *r* taken from a shuffled vector of size *m* the size of *U* is simply the binomial coefficient 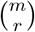:

We can then estimate *p*-values using a standard approximation (because under *H*_0_ the observed value arises from the null distribution and should therefore be included) (3) as:

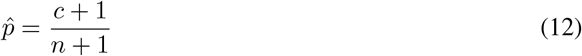

where *c* is the number of null distribution samples with a value less extreme than the observation.

Phipson and Smyth (15) give a formula for an exact *p*-value when sampling with replacement that accounts for the duplication of samples discussed above, but when *m >> r*, |*U*| grows exponentially with respect to *r* and then *E*(*n*) → *d* for realistic *d*, this correction is only useful at the smallest region sizes. Since their method involves computing a binomial cumulative probability distribution for each sample, for efficiency we simply use equation 11 to correct *n* when estimating *p*-value error by sampling with replacement.

From (9) it can be seen that the empirical significance estimation procedure is exponential in the logarithm of the minimum *p*-value obtained. Depending on the distribution of values to be tested, this means that accurately quantifying the significance of the most extreme points can become computationally infeasible (figure 3.2B and D).

Instead, we adopt an approach that involves performing a reasonably small number of permutations to generate an initial empirical distribution, then fitting theoretical distributions to the tails of the distribution. Following (10) we use the generalized Pareto distribution by default, though any distribution in the SciPy stats library (12; 14) may be specified. This approach can be used to estimate *p* values in ranges that would be completely refractory to straightforward permutation testing.

As in the vector method, we require a distinct null distribution for each considered region size. Therefore we generate an initial set of empirical distributions by repeatedly shuffling the row and column indices in parallel (such that the order of loci is randomized but the pairwise relationships between loci are preserved) and computing a null score matrix *S*_0_ for each shuffle. We generate the null distributions for each region size from the corresponding diagonals of the *S*_0_ matrices. The number of matrix shufflings is determined according to the size of the matrix such that the maximum region size tested will have at least 10,000 permuted data points.

### Implementation of empirical distribution methods

The histogram approximation method is implemented as a Python class in the `empdist` package with an interface that mirrors most of the methods of the frozen continuous variable classes in the Scipy stats module, but which also implements standard arithmetic operations between instances using the convolution approach. As of version 0.19.0 Scipy provides an `rv histogram` class with similar functionality but it does not implement the standard approximation for *p*-value reporting (which can result in values of 0) and, crucially, it does not support operations for computing expected arithmetic outcomes via convolution. Additionally, the `empdist` package provides helper functions for computing the expected distribution of the minimum and maximum of two empirical distributions. The dynamic programming methods for computing region scores are implemented in C for performance reasons.

The piecewise hybrid method is implemented as an additional Python class in the `empdist` package. The support of the distribution is divided into a left tail, central region and right tail and the cutoffs dividing the three regions are computed as the values at which an empirical *p*-value exceeds a given 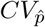 threshold (default 0.05), for the number of samples being fit (the user can also input the expected number of unique samples, as above, if it differs from the number of input samples). The central region between the tail cutoffs is modeled using the histogram-based empirical distribution described above, while the left and right tails are modeled with generalized Pareto distributions. These distributions are fit over the set of input data that is between the 0.9 quantile (0.1 for the left tail) and the most extreme confident quantile for a given 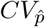 threshold. The fit is accomplished using a nested approach that speeds up the parameter search and improves its robustness compared to a naive approach. It also incorporates penalty terms to ensure that the support of the resulting distribution encloses the range of values (region scores) to be tested. The following procedure is given for the right tail; an equivalent procedure is followed for the left tail, fitting negated copies of the smallest quantiles:

A. Compute the empirical *p*-values using the fit data for 100 evenly-spaced points over the range between the 0.9 quantile and the quantile of the minimum *p*-value obtainable for the given number of sampled points at the specified CV threshold.
B. Use the Nelder-Mead algorithm (13) to find a shape parameter that maximizes the Pearson correlation of the target data values and the quantile function of a generalized Pareto distribution for the target *p*-values. The correlation score is penalized with a constant term if the support of the distribution fails to enclose the tail cutoff value and / or the maximum score to be quantified plus a variable penalty that is proportional to the difference between the support endpoint and the required support endpoint.
C. Using the optimal shape parameter discovered above, compute the corresponding location and scale parameters.
D. Use the Nelder-Mead algorithm to simultaneously optimize the shape, location and scale parameters of a generalized Pareto distribution that minimizes the squared error of the log survival function of the target data points with the log of the target empirical *p*-values. This optimization is initialized with the parameter values from the previous step.

### Determining significant sets of non-overlapping regions of proximal enrichment

Once null distributions have been generated using either the convolution of histograms method (as for vector input) or the tail approximation method (as for matrix input) for each region size, the log *p*-values are computed (either right-tailed, left-tailed or for both tails, depending on user input) for each element of *S* to generate a corresponding matrix *P* of −log *p*-values, with the *i*th diagonal corresponding to regions of size *i* + 1.

User-specified thresholds for minimum and maximum size, minimum absolute score and maximum *p*-value are then applied and matrix elements of *P* corresponding to invalid regions under these criteria are set to 0.

The next challenge is to choose, from many possible combinations, a set of non-overlapping significant regions. To do this, we seek to maximize the sum of negative log *p*-values in the set, which is equivalent to finding a minimum likelihood set with Fisher’s method of combining independent *p*-values by multiplication (6) (although nearby *p*-values in the matrix are clearly not independent). We use a dynamic programming approach to accomplish this in *𝒪* (*N* ^2^) time. If we consider the matrix *P* as the adjacency matrix of a directed graph such that *P*_*i*,*j*_ represents an edge between locus *i* and locus *j* ∀*j > i, d*(*i*) is the distance of the longest path from 0 to locus *i* and *T*_*i*_ is the set of upstream neighbors of *i*, then:

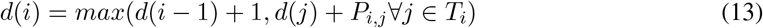

which by solving and simultaneously recording a backtrack vector we can obtain the set of non-overlapping regions with a minimum product of *p*-values.

Finally, we allow a user-specified bias parameter *α*, with a default value of 2, to which the −log *p*-values are raised prior to determination of the optimal region set. This has the effect of choosing fewer numbers of larger regions at higher values of *α* and larger numbers of smaller regions at lower values.

P-values are reported as well as adjusted *p*-values generated using the Benjamini-Hochberg method (1) (with a default false discovery rate threshold of 0.05) across all chromosomes.

## Results

### Performance of Vector PEAS

To establish a ground truth dataset we shuffled the chromosome 19 H3K27ac SPRET - C57 peak delta vector 100,000 times, generated score matrices from the shuffled vectors and used the resulting empirical null distribution of region means to compute *p*-values. Since each shuffle generates *N* − *r* samples, this resulted in approximately 5 *×* 10^8^ samples per region size, a process which took 23 hours on a server equipped with a 28-core Intel Xeon processor. For test sets we also generated 32 distinct datasets using 1,000 shuffles each of the delta vector in (each dataset was computed in approximately 13 minutes). We then fit normal distributions to the 1,000 permutations datasets and generated predicted distributions using the convolution of histograms method described above.

As can be seen in figure 3.5B, the −log_10_ *p*-values computed with the convolution approach (Vector PEAS, “convolved”) track the empirical values from 100,000 shuffle ground truth set while the normal approximations are inaccurate at small region sizes, consistent with the leptokurtic distributions there, but becomes a good estimator for large region sizes, consistent with the central limit theorem. This performance is also reflected in the maximum error computed over the range for which the 100,000 permutation set is reliable (figure 3.5C). The empirical −log_10_ *p*-values derived from the 1,000 shuffle dataset closely match the ground truth values up to the maximum value that can be computed with *CV*_*p*_ *<* 0.05 (see equation 10), but then become unstable, whereas the convolved predictions continue tracking the ground truth values up to the ground truth confidence limit, and remain on that trend even after the point where we canot make confident predictions using the the ground truth permutations. Comparing the −log_10_ *p* values for the convolved and normal approximations against the empirical −log_10_ *p* values over the confident range of scores (figure 3.5) likewise shows good agreement for the convolution method across the region sizes and poor performance of the normal approximation at small region sizes.

We used Vector PEAS to call CRDs on the on the 19 H3K27ac ChIP-seq SPRET-C57 Δ autosomal chromosome vectors with a *p*-value threshold of 0.005, an FDR threshold of 0.05, a minimum size of 3 and *α* set to 2 (figure 3.6) resulting in a set of genome-wide set of 2438 H3K27ac CRDs. We also called CRDs on the SPRET-C57 Hi-C PC1 Δ vectors (after first smoothing large-scale deviations by taking the residuals of a LOWESS regression with a 10 MB window. See figure 3.8) using the same parameters, resulting in a set of 2211 PC1 CRDs. Of the total, 72.6 % of the Δ H3K27ac CRDs overlapped at least one Δ PC1 CRD, and 59.5 % of the Δ PC1 CRDs overlapped at least one Δ H3K27ac CRD. The majority (70.3 %) of the 2044 pairwise overlap events were concordant (between Δ CRDs with the same sign, indicating inter-specific change in the same direction) (figure 3.7A). This concordance is consistent with the increased activation state of regulatory regions in A compartments. Concordant overlapping CRDs had higher absolute score values (figure 3.7B, Mann-Whitney p-values: 1.84 *×* 10^−15^ for Δ PC1 CRDs, 1.8*×*10^−3^ for Δ H3K27ac CRDs). 51 of the 2044 overlap events had a Jaccard coefficient greater than 0.75, indicating a very close correspondence of their boundaries, and 49 of these were concordant. Such a close overlap may represent a chromatin domain whose activity is altered by changes in *cis* or *trans* regulatory features between species. Alternatively, a shift in chromatin compartment membership between the species may have altered the accessibility of intra-domain binding sites that then resulted in decreases in the H3K27ac mark.

### Performance of Matrix PEAS

Analogously to the procedure used for evaluating the performance of Vector PEAS, above, we generated a ground truth set using 100,000 shuffled correlation matrices and 32 smaller datasets using 1,000 shuffled matrices. A normal distribution was fit to each of the 32 smaller datasets, as well as a piecewise tail approximation distribution as used in Matrix PEAS. The relative accuracy of the −log_10_ *p*-values computed by the various approaches compared to the ground truth set can be observed in figures 3.5B and 3.5D, with the piecewise tail approximation (TA) approach of Matrix PEAS robustly approximating the tail behavior of the empirical distributions over the range for which the ground truth data gives confident estimation. Beyond this confident assessment range (the double-crosshatched region of 3.5B), the log_10_ *p*-values maintain a reasonable trend but exhibit a high variance at larger region sizes because the very long tail must be extrapolated from a small number of data points in the central region. While this variance is undesirable, it remains preferable to alternative methods of p-value estimation that are either computationally-intractable (naive permutation) or very inaccurate (theoretical distributions estimated over the entire dataset, e.g. normal). To some extent this variance can be reduced by increasing the number of permutations used to fit the piecewise TA distribution. In contrast to the application of Vector PEAS, the normal distribution is a reasonable fit to the data at small region sizes, but becomes much worse at larger sizes. This is because of the excess of positive correlation values at large region sizes in the true correlation matrices, which create null distributions having reasonable kurtosis but skewed to the large positive tail.

**Figure 3.6:**
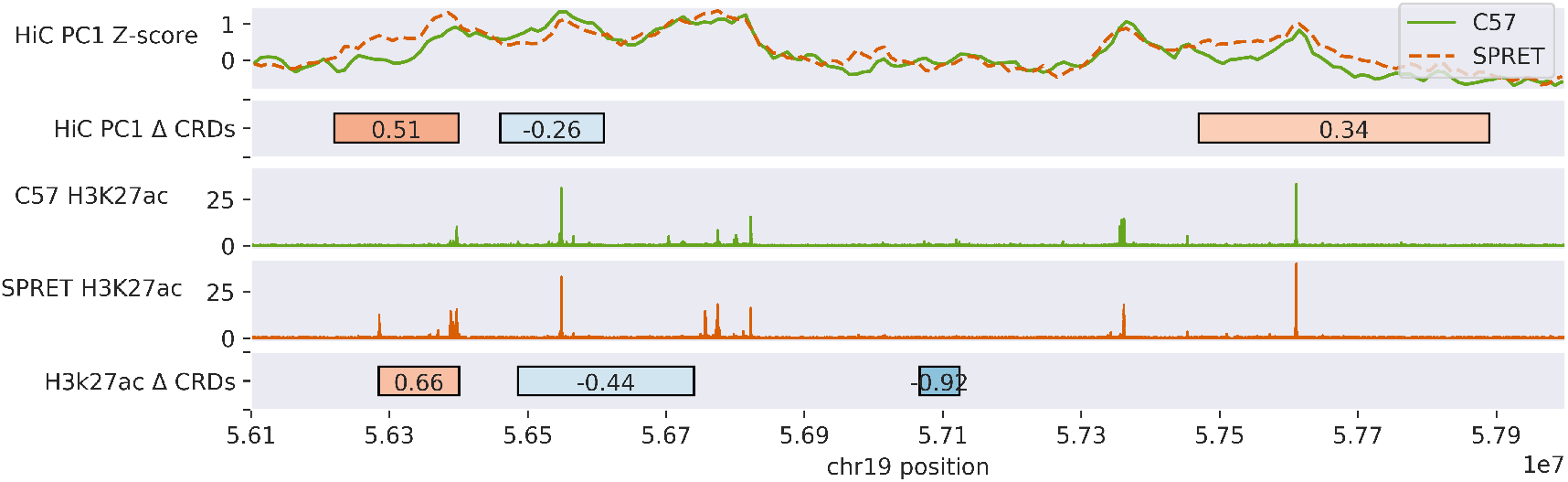
CRDs (as boxes) and input data for SPRET and C57 BMDMs over a section of chromosome 19. CRDs are called with vector PEAS from differences in log-transformed H3K27ac peak heights and on the difference in z-scores of the first principle component of the Hi-C correlation matrices. CRDs are shaded with red for regions greater in SPRET, and in blue for regions greater in C57. Corresponding normalized ChIP-seq read profiles as well as the Hi-C PC1 z-scores are shown above the CRDs.

## Discussion

The related approaches of PEAS offer a number of advantages over existing methods. First, they compute the statistical significance of candidate CRDs, thus providing a straightforward way for researchers to stratify findings and determine thresholds on the basis of confidence rather than arbitrary heuristics. By using an empirical approach it does so without requiring any assumptions about the distributions of the underlying data and easily accommodates unusual distributions, reducing upstream requirements for data normalization. By approximating a large permutation test, it can compute very small *p*-values with a reasonable computation time (*<* 1 hr on a single-core machine).

A common approach for computing significance in similar situations is to use a computationally-tractable theoretical distribution such as a Gaussian and rely on the central limit theorem to ensure a reasonable fit to the data. Indeed, the distributions of the scores of the largest regions, comprising the mean of many random variables, do approach normality. However, as we have shown, this convergence to normality is slow enough that *p*-values computed in the significant tail region will be incorrect by many orders of magnitude. Even small errors in *p*-values can have dramatic effects on the family-wise error rate when combined in multiple testing contexts (15). Methods of combining *p*-values are similarly dependent on accurate estimations.

**Figure 3.7:**
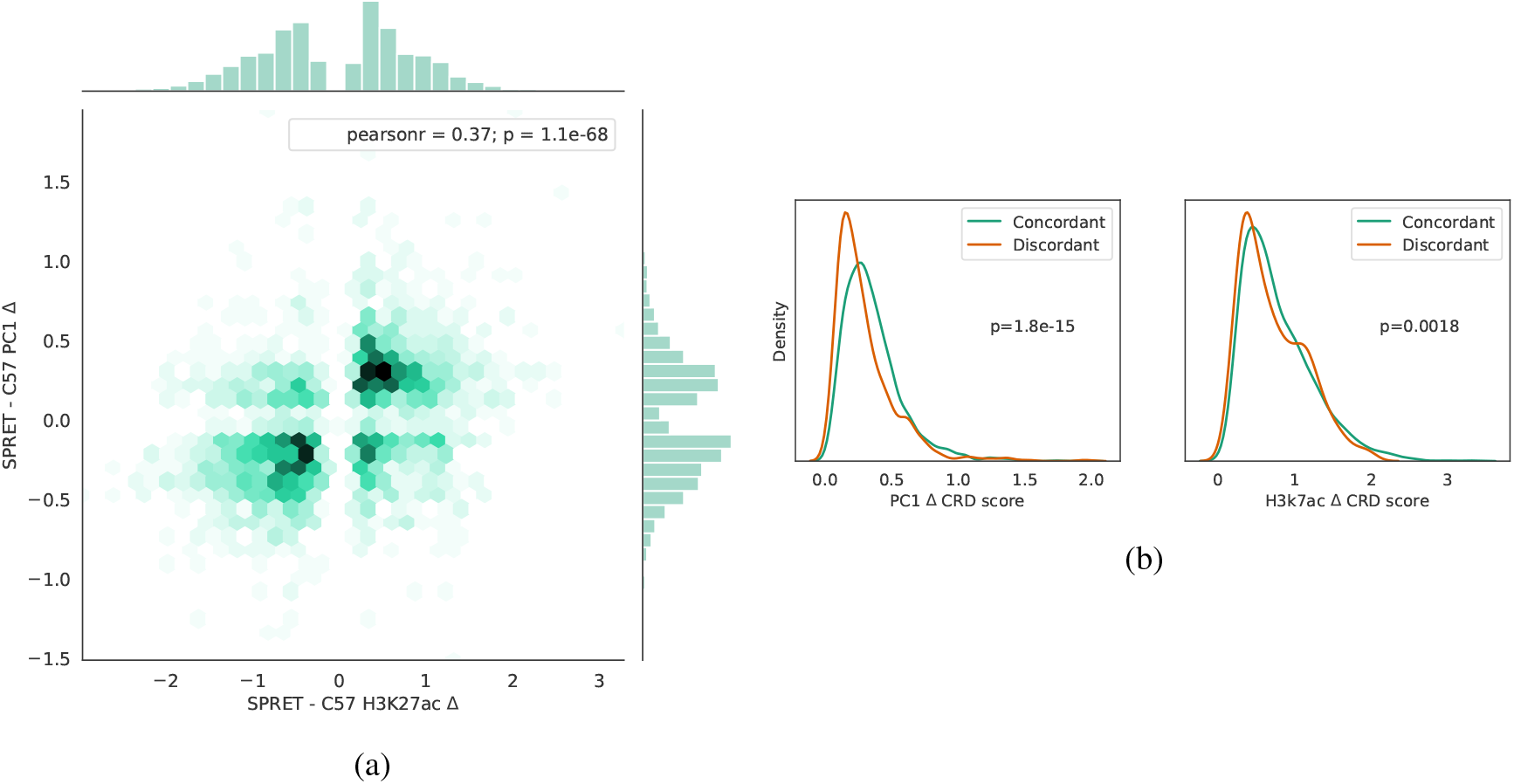
(A) Relative density of H3K27ac CRD scores vs. HiC PC1 CRD scores for overlapping CRDs genomewide. (B) Differences in score distributions for concordant vs. discordant overlapping H3K27ac and PC1 CRDs genomewide.

One shortcoming of the PEAS methods is that, unlike methods that rely on an initial hierarchical clustering step, PEAS is not able to detect *trans* regions of coordinated change on non-contiguous regions of a chromosome or across chromosomes. However, by focusing solely on *cis* regions we greatly reduce the vulnerability to false positives that would occur when searching for *trans* interactions using correlations between a handful of samples.

Although the methods shown have been demonstrated using a peak-based approach, they work equally well when dealing with read counts over equal-sized bins tiled evenly across the genome. Such binned approaches have the advantage of implicitly taking genomic distance into account, unlike peak-based methods.

In addition to offering a robust solution to the original motivating problem of identifying CRDs, the methods are generalizable to any ordered observation vector or matrix and may be useful in other contexts, e.g. identifying copy number variants from contiguous regions of deviation in read counts along a chromosome.

**Figure 3.8:**
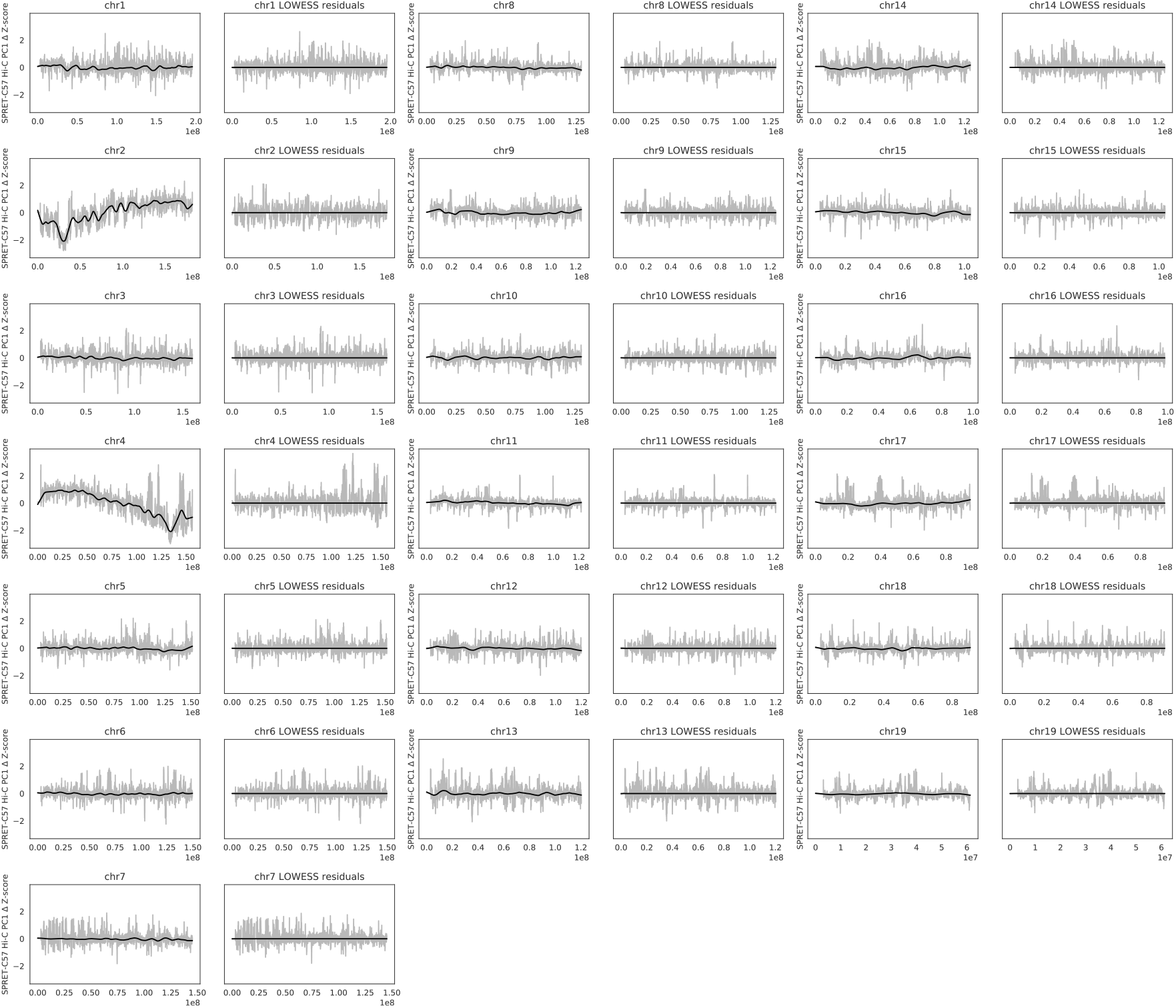
SPRET-C57 Hi-C PC1 Δ vectors for each chromosome alongside the corresponding residuals of a 10 MB window LOWESS regression.

## Conclusion

We present a software package that implementation two different methods for efficiently estimating empirical distributions to allow accurate computation of *p*-values for data quantiles that would be infeasible to assess using naive permutation testing: using histogram convolution to accurately predict the outcome of arithmetic operations on independent empirical distributions and using tail approximations to extrapolate the results of a small number of permutationsin cases where independence does not hold. We present a software package, PEAS, that applies these methods to the general problem of computing the significance of enriched or depleted regions along an ordered vector or matrix, respectively, as well as to the specific case of finding CRDs in genomic data using small sample sizes.

The PEAS method is the first, to our knowledge, that provides the ability to delineate both vector and matrix CRDs using their likelihood under a simple null model free of assumptions about the distribution of individual region scores and without relying on arbitrary heuristics. It offers fast performance and flexibility with the ability to threshold on multiple metrics (mean score, *p*-value, minimum and maximum size) simultaneously and allows the user to override the default settings of most parameters.

## Acknowledgements

Chapter 3, in part, is currently being prepared for submission for publication of the material. Skola, Dylan; Glass, Christopher. The dissertation author was the primary investigator and author of this paper.

## Bibliography

[1] Y. Benjamini and Y. Hochberg. Controlling the False Discovery Rate: A Practical and Powerful Approach to Multiple Testing. Journal of the Royal Statistical Society. Series B (Methodological), 57(1):289–300, 1995.

[2] J. D. Buenrostro, P. G. Giresi, L. C. Zaba, H. Y. Chang, and W. J. Greenleaf. Transposition of native chromatin for fast and sensitive epigenomic profiling of open chromatin, DNA-binding proteins and nucleosome position. Nature Methods, 10(12):1213–1218, Dec. 2013.

[3] A. C. Davison and D. V. Hinkley. Bootstrap methods and their application. Cambridge University Press, Cambridge; New York, NY, USA, 1997. ISBN 978-0-521-57391-7 978-0-521-57471-6.

[4] O. Delaneau, M. Zazhytska, C. Borel, C. Howald, S. Kumar, H. Ongen, K. Popadin, D. Marbach, G. Ambrosini, D. Bielser, D. Hacker, L. Romano-Palumbo, P. Ribaux, M. Wiederkehr, E. Falconnet, P. Bucher, S. Bergmann, S. Antonarakis, A. Reymond, and E. Dermitzakis. Intra- and inter-chromosomal chromatin interactions mediate genetic effects on regulatory networks. bioRxiv, page 171694, Aug. 2017.

[5] J. R. Dixon, S. Selvaraj, F. Yue, A. Kim, Y. Li, Y. Shen, M. Hu, J. S. Liu, and B. Ren. Topological domains in mammalian genomes identified by analysis of chromatin interactions. Nature, 485 (7398):376–380, Apr. 2012.

[6] R. A. Fisher. 224a: Answer to Question 14 on Combining independent tests of significance. 1948.

[7] R. E. Gate, C. S. Cheng, A. P. Aiden, A. Siba, M. Tabaka, D. Lituiev, I. Machol, M. G. Gordon, M. Subramaniam, M. Shamim, K. L. Hougen, I. Wortman, S.-C. Huang, N. C. Durand, T. Feng, P. L. D. Jager, H. Y. Chang, E. L. Aiden, C. Benoist, M. A. Beer, C. J. Ye, and A. Regev. Genetic determinants of co-accessible chromatin regions in activated T cells across humans. Nature Genetics, 50(8):1140, Aug. 2018.

[8] S. Heinz, C. Benner, N. Spann, E. Bertolino, Y. C. Lin, P. Laslo, J. X. Cheng, C. Murre, H. Singh, and C. K. Glass. Simple combinations of lineage-determining transcription factors prime cisregulatory elements required for macrophage and B cell identities. Molecular Cell, 38(4):576–589, May 2010.

[9] D. S. Johnson, A. Mortazavi, R. M. Myers, and B. Wold. Genome-Wide Mapping of in Vivo Protein-DNA Interactions. Science, 316(5830):1497–1502, June 2007.

[10] T. A. Knijnenburg, L. F. A. Wessels, M. J. T. Reinders, and I. Shmulevich. Fewer permutations, more accurate P-values. Bioinformatics, 25(12):i161–i168, June 2009.

[11] V. M. Link, S. H. Duttke, H. B. Chun, I. R. Holtman, E. Westin, M. A. Hoeksema, Y. Abe, D. Skola, C. E. Romanoski, J. Tao, G. J. Fonseca, T. D. Troutman, N. J. Spann, T. Strid, M. Sakai, M. Yu, R. Hu, R. Fang, D. Metzler, B. Ren, and C. K. Glass. Analysis of Genetically Diverse Macrophages Reveals Local and Domain-wide Mechanisms that Control Transcription Factor Binding and Function. Cell, 173(7):1796–1809.e17, June 2018.

[12] K. J. Millman and M. Aivazis. Python for Scientists and Engineers. Computing in Science Engineering, 13(2):9–12, Mar. 2011.

[13] J. A. Nelder and R. Mead. A Simplex Method for Function Minimization. The Computer Journal, 7(4):308–313, Jan. 1965.

[14] T. E. Oliphant. Python for Scientific Computing. Computing in Science Engineering, 9(3):10–20, May 2007.

[15] B. Phipson and G. K. Smyth. Permutation P-values Should Never Be Zero: Calculating Exact Pvalues When Permutations Are Randomly Drawn. Statistical Applications in Genetics and Molecular Biology, 9(1), 2010.

[16] H. A. Pliner, J. S. Packer, J. L. McFaline-Figueroa, D. A. Cusanovich, R. M. Daza, D. Aghamirzaie, S. Srivatsan, X. Qiu, D. Jackson, A. Minkina, A. C. Adey, F. J. Steemers, J. Shendure, and C. Trapnell. Cicero Predicts cis-Regulatory DNA Interactions from Single-Cell Chromatin Accessibility Data. Molecular Cell, 71(5):858–871.e8, Sept. 2018.

[17] A. R. Quinlan and I. M. Hall. BEDTools: a flexible suite of utilities for comparing genomic features. Bioinformatics, 26(6):841–842, Mar. 2010.

[18] S. M. Waszak, O. Delaneau, A. R. Gschwind, H. Kilpinen, S. K. Raghav, R. M. Witwicki, A. Orioli, M. Wiederkehr, N. I. Panousis, A. Yurovsky, L. Romano-Palumbo, A. Planchon, D. Bielser, A. Padioleau, G. Udin, S. Thurnheer, D. Hacker, N. Hernandez, A. Reymond, B. Deplancke, and E. T. Dermitzakis. Population Variation and Genetic Control of Modular Chromatin Architecture in Humans. Cell, 162(5):1039–1050, Aug. 2015.

